# Rhinovirus Infection of Human Airway Epithelial Cells Stimulates Neutrophil Superoxide

**DOI:** 10.64898/2026.10.05.756757

**Authors:** Jo L. Wilson, R. Brockman-Schneider, Terry D. Juang, Gretta A. Kennedy, Mark Devries, Yury A. Bochkov, Sheena C. Kerr, James E. Gern

**Author notes:** Corresponding Author Jo L. Wilson, MD, Department of Pediatrics, University of Wisconsin–Madison, Madison, WI, USA.

## Abstract

**Background:** Neutrophils migrate to the respiratory epithelium during rhinovirus (RV) infection, but whether RV directly activates neutrophils or whether activation arises from epithelial responses remains unclear.

**Objective:** To determine whether RV directly activates human neutrophils or indirectly activates them through RV-infected airway epithelial cells.

**Methods:** Human neutrophils were exposed to purified RV, and activation was assessed by myeloperoxidase activity, CD11b/CD66b expression, and reactive oxygen species (ROS) production. Neutrophils were co-cultured with RV-infected air–liquid interface (ALI) bronchial epithelium evaluated for effects on epithelial integrity. At 48-72 hours post-infection, we used live-cell imaging to measure ROS responses and neutrophil-tracking assays to measure migration velocity and ROS responses to basolateral supernatants, clarified apical washes, and apical debris from RV-infected epithelial cells. Plaque assays were used to assess for RV infectivity after incubation with apical neutrophils.

**Results:** RV did not directly activate neutrophils. Adding neutrophils to the apical surface of airway epithelial cells increased neutrophil ROS production at 48-72 hrs post-infection but did not alter epithelial cytotoxicity. Conversely, basolateral supernatants from infected cells enhanced neutrophil migration but not ROS, consistent with cytokine-mediated priming. Clarified apical washes had no effect on migration or ROS. By contrast, apical debris from infected cultures triggered a robust ROS response. Basolateral fractions from medium control- or RV-infected cultures did not induce ROS. Infectivity is not impacted by apical neutrophil presence.

**Conclusion:** RV does not directly activate neutrophils. Instead, RV-infected epithelial cells release basolateral factors that enhance neutrophil migration, while apical debris from infected cells drives ROS activation. These findings indicate that epithelial injury, not RV itself, is the dominant trigger of neutrophil activation during infection.

**Graphical Abstract:** Rhinovirus (RV) alone does not directly activate human neutrophils (no change in MPO, CD66b, CD11b, or reactive oxygen species (ROS). In contrast, RV-infected airway epithelial cells activate neutrophils. Basolateral cytokines enhance neutrophil migration, while RV-infected epithelial cell debris triggers neutrophil ROS production. Colors denote key components of the model: epithelial cells (pink/tan and orange), neutrophils (pink), rhinovirus virions (dark blue), epithelial-derived cytokines (light blue), and reactive oxygen species (red).

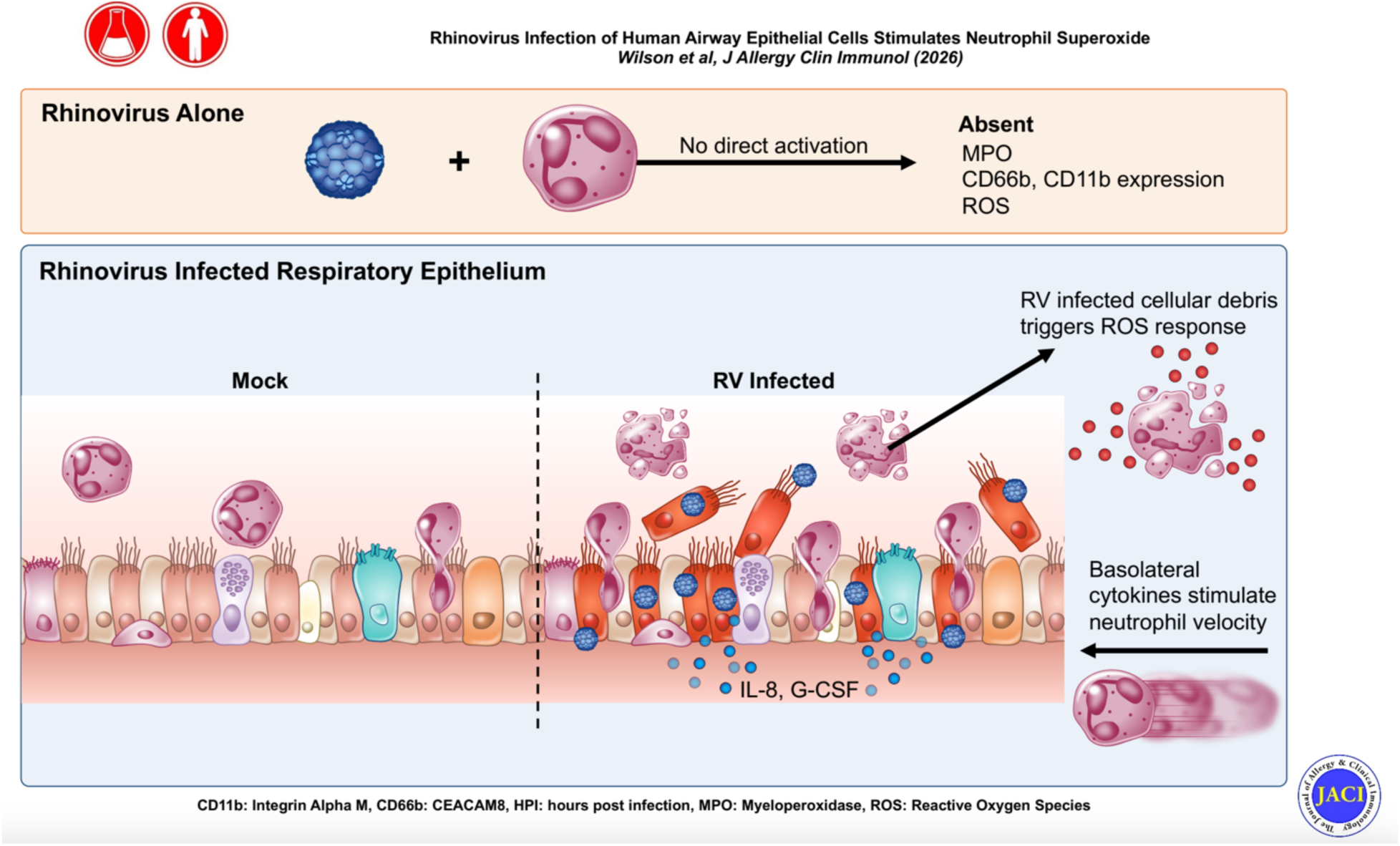

## Introduction

Rhinoviruses (RV) are significant human pathogens. While widely recognized for their contribution to over 50% of common colds, RVs also contribute substantially to morbidity in select patient populations.(1) Respiratory viral infections account for approximately 50% of adult and 80% of pediatric asthma exacerbations, with RV contributing to 60-70% of virus induced exacerbations.(2) In the case of infantile bronchiolitis, 20-30% are secondary to RV and approximately 15% of chronic obstructive pulmonary disease (COPD) exacerbations are attributed to acute RV infection.(3–6) Therefore, RV represents a pathogen responsible for significant morbidity and healthcare utilization.

RVs are members of the genus *Enterovirus* and the family *Picornaviridae*. Within this family, there are three RV species (RV-A, RV-B and RV-C) with approximately 170 types that have limited serologic cross-protection.(7) RVs infect human epithelial cells through binding to intercellular adhesion molecule (ICAM-1 - RV-A and RV-B), low-density lipoprotein receptor (LDLR - RV-A), or cadherin-related family member 3 (CDHR3 - RV-C) receptors. CDHR3 expression is limited to ciliated cells in the airway, which are the exclusive host cell for RV-C infections.(8)

Neutrophils are the most abundant white blood cells and are critical in innate immunity. Neutrophils exert antimicrobial activity through several conserved effector functions including phagocytosis, production of superoxide or reactive oxygen species (ROS), degranulation of proteolytic enzymes and formation of neutrophil extracellular traps.(9) In the lung, neutrophils may be present in a steady state in lung tissue that allows for quick infiltration into airways or alveoli upon pathogen exposure.(10) Infections with respiratory viruses, including RV, stimulate neutrophil recruitment to the airway surface.(11–13) Recent mouse models of asthma show that blocking neutrophil recruitment-either through inhibition of C-X-C motif chemokine receptor 2 (CXCR2) or by interrupting interleukin-33 (IL-33)–driven inflammatory signaling-significantly reduces downstream cytokine production and airway hyperresponsiveness that depend on neutrophil influx to the airway.(14–16) However, how RV-induced neutrophil recruitment relates to their activation and clinical outcomes, including asthma exacerbations, is not well understood.

To address this research gap, we utilized human co-culture models of air-liquid (ALI) interface differentiated bronchial epithelial cells and purified human neutrophils to determine mechanisms of RV-induced activation and neutrophil effects on viral infection. We specifically addressed whether RV infections directly affect neutrophil activation or indirectly activate neutrophils through effects on airway epithelial cells. In addition, we tested whether recruited neutrophils disturb epithelial cell viability or barrier integrity. We also assessed whether incubation with neutrophils may impact RV infectivity.

## Methods

### Cell sources and processing

Human lungs rejected for transplant were obtained from the Organ Procurement Organization at UW Health under a protocol reviewed by the University of Wisconsin Institutional Review Board and determined not to constitute human subject’s research. Donor lungs were initially procured for clinical transplantation, therefore represented healthy lung donors, however subsequently deemed unsuitable for transplant. The most common reasons for non-utilization were logistical considerations, including timing and recipient-related factors.

Bronchial and tracheal epithelial cells were isolated from excised large airways by enzymatic digestion with pronase (Roche) and DNase (Sigma) at 4 °C. Fibroblasts were removed by differential adhesion to tissue-culture plastic, and epithelial cells were cryopreserved in liquid nitrogen until use. To establish ALI cultures, 500,000 to 750,000 thawed primary epithelial cells were expanded in CellBind™ T-75 flasks using PneumaCult-Ex Plus™ medium (STEMCELL Technologies) supplemented with 0.1% (vol/vol) gentamicin (Sigma) and 0.1% (vol/vol) fluconazole (Novaplus) until 60 – 70% confluent. Cells were then trypsinized and seeded onto collagen-coated (0.4 uM pore) transwell membranes (Corning Costar 3460 or 3470) at a density of 100,000 per cm^2^. Once monolayers reached confluence (typically within ∼2 days), apical medium was removed to initiate airlifting, and cultures were differentiated for at least 24 days in PneumaCult-ALI™ medium (STEMCELL Technologies), supplemented with 0.1% (vol/vol) gentamicin and 0.1% (vol/vol) fluconazole in the basal chamber. The presence of motile cilia was required to consider the culture fully differentiated.

Human neutrophils were isolated in bulk from venous blood obtained via venipuncture from healthy adult donors using the EasySep™ Direct Human Neutrophil Isolation Kit (STEMCELL Technologies, catalog #19666), following the manufacturer’s instructions. All blood samples were collected under informed consent and in compliance with institutional guidelines approved by the University of Wisconsin–Madison Institutional Review Board. To avoid inadvertent neutrophil activation, we maintained a consistent room temperature throughout isolation and handling, used polypropylene tubes (avoiding polystyrene), and used gentle pipetting techniques. Isolated neutrophils were suspended in HBSS⁺ (Hank’s Balanced Salt Solution with calcium and magnesium, 0.1% Human Serum Albumin, 20 mM HEPES). Trypan blue exclusion was used to eliminate dead cells from neutrophil counts. Donors were self-reported healthy with no chronic respiratory conditions, no evidence of acute infection, and no use of chronic immunosuppression that would impact neutrophil function. There were 6 donors (4 females and 2 males) aged 39-70.R

### Viral Infection of epithelial cultures

Rhinoviruses (RVs) were produced from infectious cDNA clones in human embryonic lung fibroblasts (WisL cells) by reverse genetics as previously described.(17–19) Viruses were purified and concentrated by ultracentrifugation (100,000 × g, 4 h, 10 °C) through a 30% sucrose cushion and resuspended in phosphate-buffered saline containing 0.01% BSA. ALI human bronchial epithelial cultures (hBEC) were infected by first removing mucus with two PBS washes, followed by inoculation of the apical surface with 100 µL (12-well insert) or 50 µL (24-well insert) of prepared virus at the desired concentrations (10³ to 10⁶ PFUe). Viral attachment was allowed to occur for 2 hours at 34 °C, 5% CO₂. The viral inoculum was then removed, the apical surfaces were washed three times with PBS, and cell lysates in RLT lysis buffer (Qiagen) were collected to measure viral attachment. Cultures were returned to an incubator (34 °C, 5% CO₂), and viral replication was allowed to proceed for the timing appropriate to individual experiments (RV Replication -15 hrs, TEER and LDH -48 hrs, ROS, 24-72 hours), at which point viral replication samples were collected. The cell lysates were harvested with RLT Lysis Buffer (Qiagen), homogenized (QIAshredder, Qiagen), and frozen at −80 °C.

### Quantitative PCR

Total RNA was isolated from cell lysates (RNeasy Mini Kits, Qiagen), and cDNA was generated (TaqMan Reverse Transcription Reagents, Applied Biosystems) using virus-specific reverse primer R848. RV RNA was quantified by real-time PCR using a standard curve and specific primers (RV1A16, R848) as previously described.(17) Negative controls lacking RNA or cDNA were included in all experiments to confirm the absence of cross-contamination.

### Neutrophil MPO production

Neutrophils (1 × 10⁶ in 50 µL) were incubated with RV (5 × 10⁶ PFUe in 50 µL) or mock control in a 96-well plate for 4 hours at RT in the dark. After incubation, 25 µL of co-incubation supernatants were analyzed in duplicate using a commercial kit (Neutrophil Myeloperoxidase Activity Assay Kit, Cayman Chemical). Absorbance at 650 nm was measured on a BioTek plate reader. *Moraxella catarrhalis* (strain MC14) was isolated by the University of Wisconsin Clinical Pathology Laboratory from a clinical specimen. Neutrophils (4 × 10⁶ in 50 µL) were co-incubated (4 hours, 37 °C, 5% CO₂) with bacteria (3 × 10⁷ CFU in 50 µL) in a 96-well plate. Co-incubation supernatants were analyzed for MPO as described above. Data represents 6 independent biologic replicates for RV studies, 3 donors for Moraxella controls.

### Flow cytometry for activation markers

Neutrophils (1 × 10⁶ in 100 µL HBSS⁺) were co-incubated (4 hours at 34 °C, 5% CO₂) with RV-C15 (5 µL of 2 × 10⁶ PFUe/µL, MOI = 10) or medium alone. Neutrophils were collected, centrifuged (400 × g, 3 minutes), and resuspended in 200 µL Cell Staining Buffer (BioLegend). Cells were stained (15 minutes in the dark) with 0.5 µL of a viability dye (Zombie NIR, BioLegend) and 5 µL of a 1:10 dilution of Apotracker Green (BioLegend). Cells were centrifuged at 400 × g for 3 minutes, washed with 750 µL Cell Staining Buffer, and centrifuged again. Pellets were resuspended in 200 µL of paraformaldehyde solution (4% in PBS; Santa Cruz) for fixation and incubated at 4 °C for 30 minutes. Samples were centrifuged (400 × g, 5 minutes) and resuspended in 200 µL Cell Staining Buffer. Fc receptors were blocked by incubating (10 minutes, RT) the 200 µL samples with 10 µL of Human TruStain FcX (BioLegend), followed by centrifugation (400 × g, 5 minutes). Pellets were resuspended in 200 µL Cell Staining Buffer. Cells were then stained by adding 5 µL each of the following antibodies (BioLegend) to the 200 µL cell suspension and incubating for 20 minutes at RT in the dark: anti-CD16 - clone 3G8) Biolegend #302008), anti-CD11b (clone M1/70 - Biolegend #101235), anti-CD66b (clone G10F5 - Biolegend #305109). Cells were pelleted (400 × g, 3 minutes), washed once with 500 µL Cell Staining Buffer, repelleted (400 × g, 5 minutes), and resuspended in 750 µL Cell Staining Buffer.

Cell suspensions were filtered through strainer-cap tubes (Falcon) with a pulse spin (500 × g, 1 minute). Flow cytometry was performed on an Attune NxT (Thermo Fisher) in the University of Wisconsin Carbone Cancer Center Flow Cytometry Laboratory (Cancer Center Support Grant P30 CA014520), and data were analyzed using FlowJo (v10.9.0). Single-stain controls, unstained controls, and fluorescence-minus-one (FMO) controls were included for all experiments and used for appropriate gating. Gating strategy for all samples included exclusion of debris, FSC-A vs FSC-H for singlets and exlcusion of dead cells (Zombie – gate). PMA stimulation (5ng/mL) was used in single stain controls to generate a bright positive signal for compensation for each fluorophore. For both controls PMA triggered a 10-fold increase in fluorescence signal for both CD11b and CD66b. Median fluorescent intensity of each gate was calculated and compared for three independent neutrophil donors comparing RV-A, RV-B and RV-C.

### Measures of epithelial integrity

Differentiated hBEC cultures grown in 12-well plates were infected (48 hours, 34 °C, 5% CO₂) with RV-A16 (1 × 10⁷ PFUe). Neutrophils suspended in 100 µL HBSS⁺ were then added to the apical surfaces of the RV-infected cells (4 hours, 37 °C, 5% CO₂). Transepithelial electrical resistance (TEER) was measured at room temperature using an EVOM2 epithelial voltohmmeter (World Precision Instruments) with an STX2 chopstick electrode, which was rinsed between wells. Basolateral culture media were collected for measurement of lactate dehydrogenase (LDH) using the CytoTox 96® Non-Radioactive Cytotoxicity Assay (Promega). Absorbance at 490 nm was read on a BioTek plate reader. Three independent neutrophil donors were used as appropriate biological replicates.

### Neutrophil ROS production

hBEC cultures were infected apically with RV-C15 (1 × 10⁶ PFUe) for 24, 48, or 72 hours prior to neutrophil addition. For each experiment, appropriate viral replication was confirmed using qPCR at both attachment and 24 HPI (Table S1). Neutrophils (1 × 10⁶ per well) were labeled with CellROX™ Deep Red Reagent according to the manufacturer’s instructions and applied to the apical surface of 24-well Transwell inserts. Live imaging was performed every 30 minutes for 12 hours using a Nikon Eclipse Ti fluorescent microscope integrated with a Pathology Devices live-cell imaging system. Imaging was conducted under controlled temperature and CO₂ conditions (37 °C, 5% CO₂) where indicated. Fluorescence was quantified in FIJI as mean fluorescence intensity (MFI) per high-power field (10X); four fields per well were imaged and averaged. MFI values were normalized to baseline fluorescence for each well.

We used non-linear regression models to evaluate changes in fluorescence over time (GraphPad Prism v10.1.1). A first-order model was used for all comparisons and fitted by least-squares regression. Group differences were assessed using an extra sum-of-squares F-test comparing model parameters B₀ and B₁. Analyses were restricted to the 200–800-minute interval following neutrophil addition. Model parameters and 95% confidence intervals are reported in Table S2.

### Neutrophil Microfluidics

Wash samples were collected by adding 200 mL of warmed PC-ALI minus medium to the apical surface of each differentiated cell layer, allowing the wash to soak for 10 minutes before transferring to microfuge tubes. In addition, the medium (supernatant) from each outer compartment of the culture wells was collected into separate tubes. Both sets of samples were centrifuged at 10,000xg for 5 minutes to separate mucus and debris from the liquid portion. Clarified aliquots of wash and supernatant samples were then removed, leaving the pelleted debris fraction in a final aliquot.

Microfluidic devices were placed into IBIDI μ-Slide 1 Well chambers (Cat. No. 82106) for the micro blood analysis technology (μBAT)(20, 21)(20, 21)-Each μBAT device contains sixteen independent channels, enabling technical triplicates for each experimental condition and multiplexed readouts. Prior to coating, the plate containing the μBAT device was placed on ice to maintain a low temperature during surface functionalization. P-selectin (R&D Systems, 50 μg/mL) was introduced into each channel (5 μL per channel) and incubated overnight at 4 °C to allow channel coating. Following neutrophil isolation as previously described, neutrophils were introduced directly into the μBAT channels for capture on the P-selectin–coated surfaces, which had been prefilled with EGM-2 MV containing dihydrorhodamine 123 (DHR123) and Hoechst stain. Neutrophils were allowed to adhere for 10 minutes, after which non-adherent cells were removed by washing the channels three times with 6 μL of EGM-2 MV. During these wash steps, the media contained dihydrorhodamine 123 (DHR123) and Hoechst stain to label cells for subsequent ROS and nuclear visualization. After the initial 10-minute capture period, media corresponding to the different experimental conditions was introduced into the channels; control conditions consisted of culture media containing only the staining dyes (DHR123 and Hoechst). PMA (50 nM, Abcam) was used as a stimulated positive control for reactive oxygen species (ROS) generation. Neutrophil motility was quantified by tracking cell nuclei using the Hoechst (blue) fluorescence channel, while ROS intensity was quantified from the green fluorescence channel. Time-lapse fluorescence imaging was performed using a Nikon Ti Eclipse inverted epifluorescence microscope equipped with a motorized XY stage and an environmental chamber providing gas and humidity control at 37 °C for live-cell imaging. Imaging was conducted using a 20× air objective with bright-field illumination and two excitation wavelengths (390 nm and 485 nm) corresponding to Hoechst and DHR123 fluorescence, respectively. μBAT devices were imaged for up to 6 hours at 3-minute intervals, with each experimental condition run in triplicate within a single device unless otherwise noted. Image analysis and cell tracking were performed using FIJI (ImageJ) with the TrackMate plugin(20) to quantify neutrophil migration. Cell Velocity was calculated using the IBIDI Chemotaxis and Migration Tool (IBIDI, Gräfelfing, Germany).(21, 22)

### Neutrophil and RV-A16 Plaque Assays

hBECs were infected with RV-A16 (5 × 10^7^ PFUe) for 72 hours and then incubated with neutrophils (1×10^6^/well) obtained as described above for 12 hours to assess effect of ROS on infectivity. After 12 hours, apical washes were collected, sonicated, centrifuged to remove cellular debris and frozen at –80°C. Prior to plaque assay, three freeze-thaws occurred and the pellet was resuspended in the same volume as the supernatant. Samples were then plated on HeLa-E8 cell monolayers (95-100% confluence) as follows: Growth medium was aspirated from the HeLa-e8 cells, and the cells were washed with warm PBS+. Following aspiration of PBS+, the cells were infected with 175 uL of each inoculant. The cells incubated with viral inoculum at room temperature for one hour on a rocking plate. Following incubation, an agar overlay consisting of 50% P6 medium (Eagles MEM powder (Gibco 41500-034), 2.2 g sodium bicarbonate, 8.1g MgCl2*6H2O in 450ml ddH2O, 13.3 ml of 7.5% bovine serum albumin Fraction V (Gibco 15260-037), pH adjusted to 7.0) and 50% 1.6% agarose was prepared and added to the cell monolayers. A liquid overlay was added after agar solidified (50% P6 medium, 48% ddH2O, 1% GOP (100X stock-2.92g L-glutamine, 1.1g pyruvic acid, 2.64g oxaloacetic acid, brought to volume of 100ml, filter sterilized, and stored frozen at -20 °C), 1% D-glucose (20% w/w stock). Liquid overlay (1.5 mL) was added to each well and HelaE8 monolayers were incubated at 34°C for 72 hours. Agar and liquid overlays were removed, and the cells were fixed with 10% formalin for at least one hour. Cells were stained with 0.1% crystal violet solution (in 20% ethanol). The crystal violet solution was removed after one hour. Data represents 5 independent biologic replicates (hBEC donors).

## Results

### Coincubation of RV and human neutrophils does not activate neutrophils

To determine whether RV species can directly activate neutrophils, we tested whether incubating neutrophils with high doses (5 × 10⁶ PFUe, MOI = 5) of RV-A, RV-B, or RV-C induced degranulation via myeloperoxidase (MPO) or upregulation of neutrophil surface activation markers, CD11b and CD66b, which are associated with increased adhesion, migration and oxidative burst.(23) *M. catarrhalis*, a common airway pathobiont, strongly increased MPO production (Fig 1A). In contrast, none of the three species of RV induced MPO greater than the control samples (Fig 1B). Similarly human neutrophils incubated with RV showed no signs of direct activation via either upregulation of CD11b (Fig 1C) or CD66b expression (Fig 1D).

**Fig 1.**
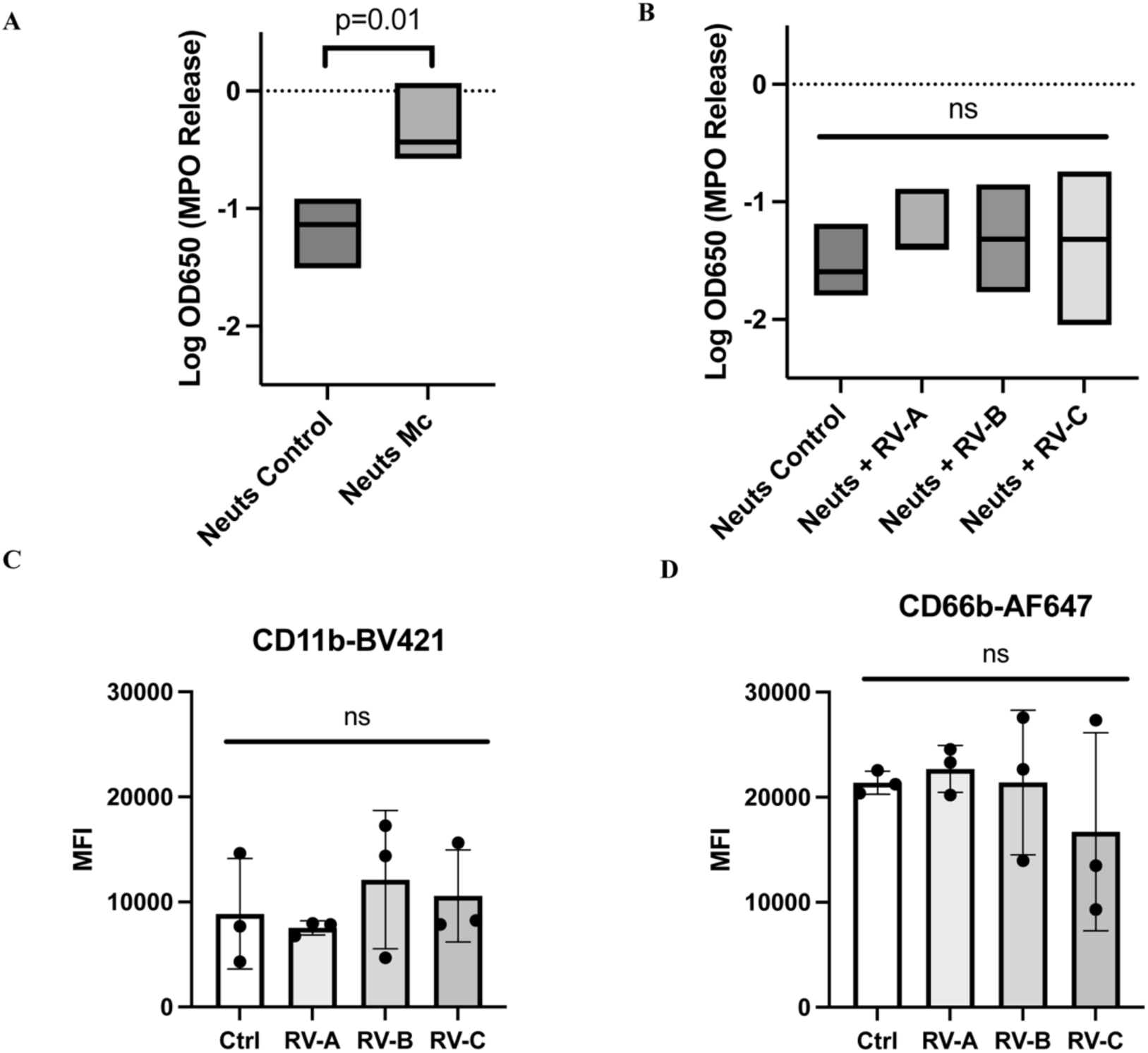
Direct neutrophil response to RV and *M. catarrhalis*. Neutrophils were incubated directly with (A) *Moraxella catarrhalis* (Mc) and (B) Rhinovirus (RV) A, B and C and MPO release was measured. *Moraxella catarrhalis* controls, n=3 donors, RV-A,B,C, n=6 donors. *Moraxella* vs. Control, p= 0.01. All RV samples compared to control were non-significant, all p > 0.31. Neutrophils were incubated directly with RV-A, RV-B and RV-C. Surface activation markers, CD11b (C) and CD66b (D) were assessed by flow cytometry. RV-A, RV-B and RV-C were all non-significant compared to Control (Ctrl), all p > 0.57.

### Neutrophils on RV-infected airway epithelium do not alter barrier integrity or cytotoxicity

To assess whether neutrophils increase RV-induced epithelial injury, neutrophils were co-cultured with RV-infected differentiated airway epithelium, and transepithelial electrical resistance (TEER) and lactate dehydrogenase (LDH) release were measured. RV infection reduced TEER relative to uninfected controls, but the addition of neutrophils did not further decrease TEER (Fig 2A). RV infection also increased LDH release to a similar degree with or without neutrophils (Fig 2B).

**Fig 2.**
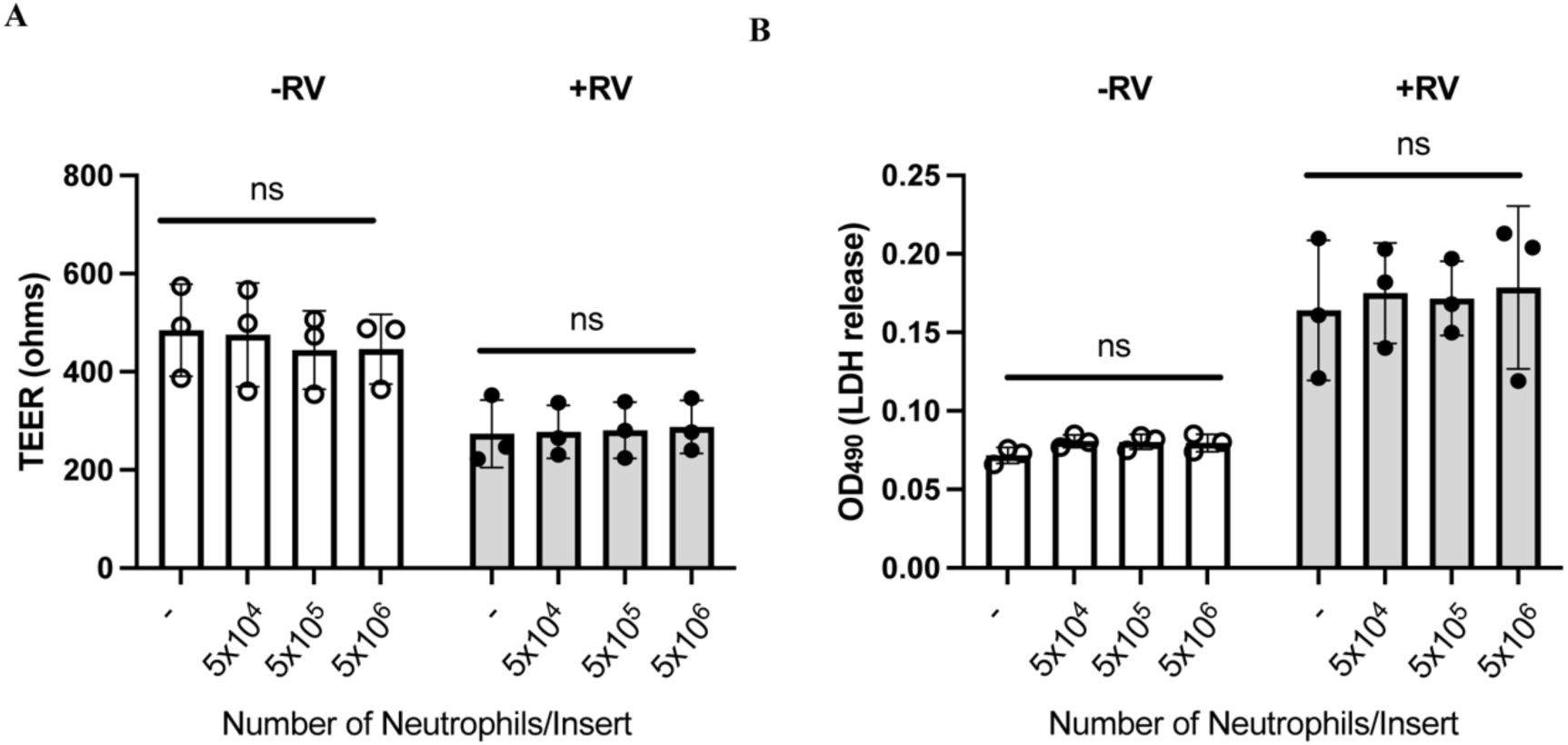
Neutrophils do not impact RV-induced changes in transepithelial resistance (TEER) or LDH release. Neutrophils were incubated on the apical surface of airway epithelial cells to assess the effect of neutrophils on the epithelial barrier. (A) TEER with media-control infection (open circles) and with RV-C infection (closed circles) and (B) LDH release with media-control infection (open circles) and with RV-C infection (closed circles). Each point represents one biological replicate. Statistical analysis performed using 2way ANOVA with multiple comparisons. Mock vs. RVC-15 p<0.0001, no significant effect of neutrophil abundance on apical surface (all p > 0.91) for both TEER and LDH.

### Neutrophils are activated by RV-infected epithelium at later stages of infection

Our data suggests that neutrophils are not directly activated in response to RV exposure, suggesting that RV infection activates neutrophils indirectly through effects on airway epithelial cells. To test this hypothesis, we infected hBEC with RV-C15, added neutrophils 24, 48, and 72 hours post-infection (HPI) and then measured neutrophil ROS production via live cell imaging at 30-minute intervals for an additional 12-hours of co-culture. At 24 HPI, introduced neutrophils exhibit a statistically significant but variable difference in ROS production compared to control infected cultures (Fig S2). Infected cultures at 48 and 72 HPI showed a consistent increase in ROS production compared to media control-infected controls (Fig 3 A-C). Notably, RV-infected hBEC without neutrophils did not increase ROS (Fig 3B).

**Fig 3.**
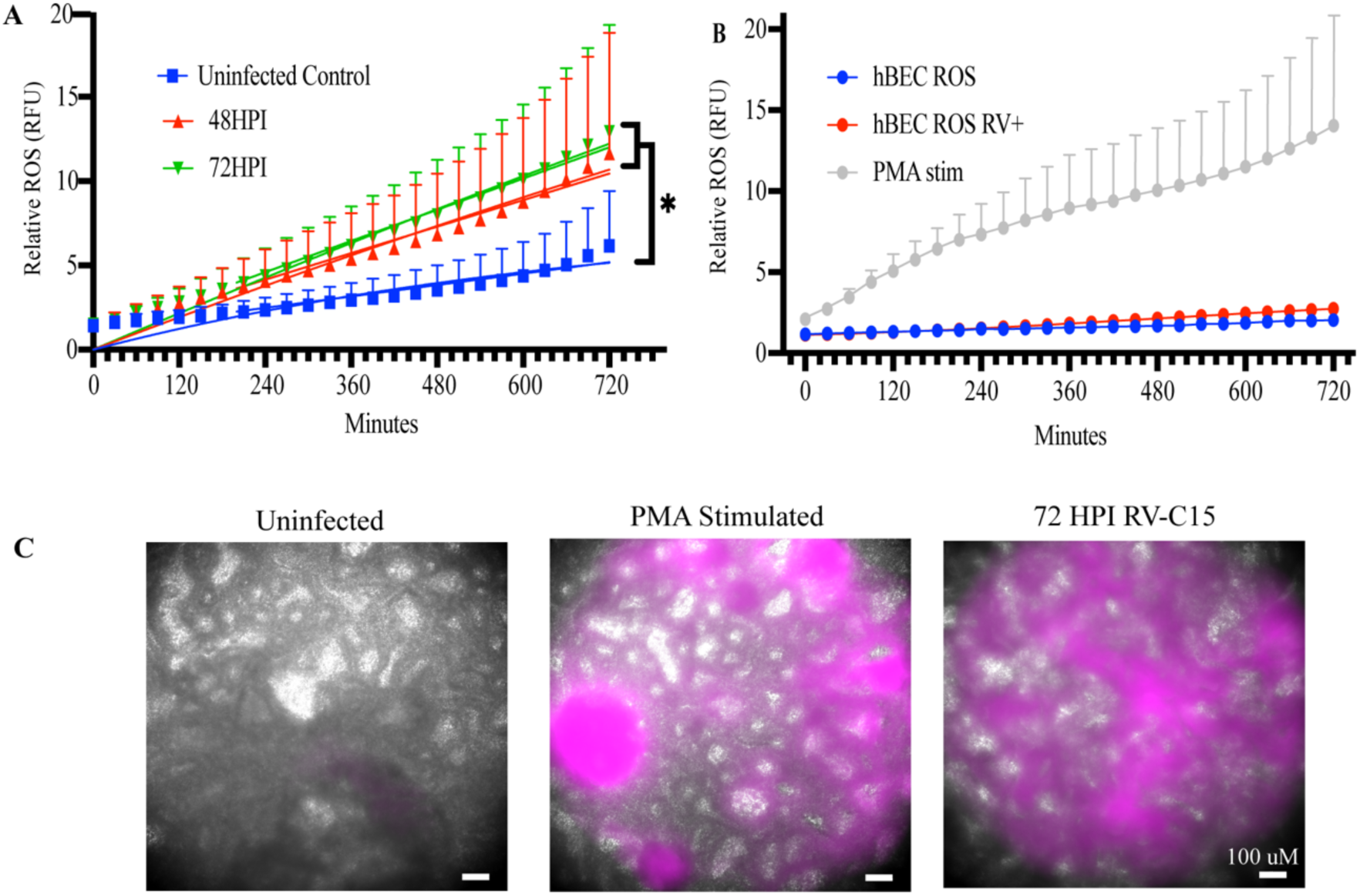
RV-C-infected epithelium induces neutrophil ROS production. (A) hBECs were inoculated with RV-C15 or medium alone and allowed to replicate for 48 or 72 hours, at which point neutrophils (1×10^6^/well) were added to the apical surfaces of the cell layers in the presence of CellROX™ Deep Red. (B) hBECs were inoculated with RV-C15 and imaged as in panel A, Neutrophils with PMA is shown to show as positive control. Epithelial cells incubated with CellROX™ to demonstrate the hBEC addition of ROS after infection are shown from cells with and without RV-C infection. (C) Representative images at 540 minutes of co-culture demonstrating visible fluorescence due to neutrophil ROS.

### Directional effects on neutrophil velocity and ROS

RV infection can induce directional (apical vs. basal) secretion of cytokines and chemokines.(24) We therefore used a microfluidic system(25) (Fig 4A) to test whether RV induced epithelial cells to directionally secrete factors regulating neutrophil chemotaxis and ROS production. We inoculated hBEC with RV-C15 or medium alone and collected both the basolateral supernatant and apical washes from the apical surface, and the latter were fractionated to evaluate effects of the liquid and cellular components. The RV-infected basolateral supernatant increased neutrophil velocity compared to media from uninfected hBECs (Fig 4B), while the RV-C infected apical washes did not. PMA decreased neutrophil velocity compared to control (Fig 4B). Basolateral supernatants from infected cells did not induce ROS production (Fig S3), while apical wash that included debris from apically shed virally infected cells markedly increased neutrophils ROS production (Fig 4 C). Notably, clarified apical washes did not increase ROS (Fig 4C).

**Fig 4.**
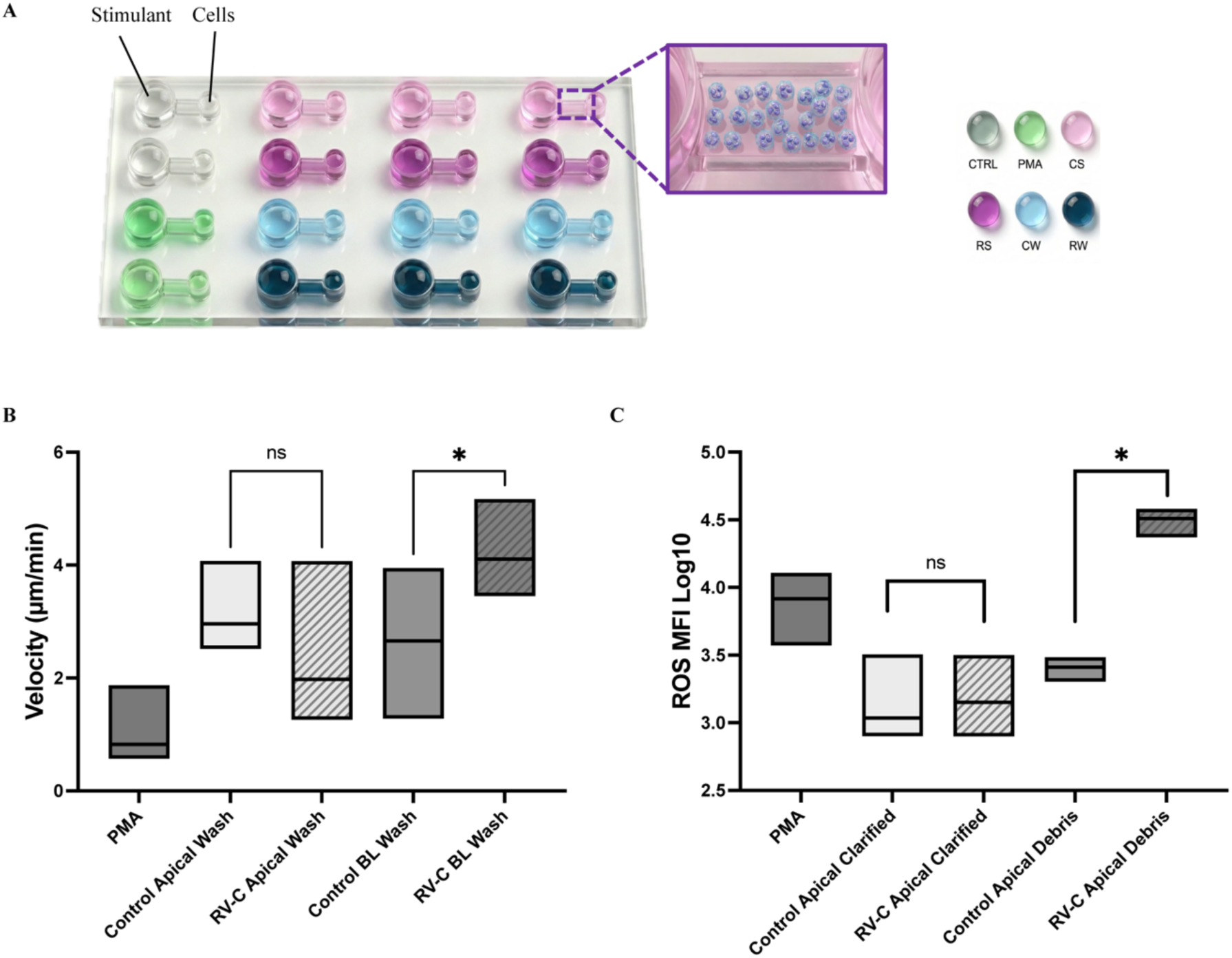
RV-infected basolateral supernatant induces neutrophil velocity, while infected apical debris induces ROS production. hBECs were incubated with RV-C15 or medium alone. Apical washes and basolateral supernatants were collected. Samples from control and RV infected samples were evaluated for their effects on freshly isolated neutrophils using microfluidic microscopy assays (A). Outcomes included neutrophil velocity (μm/min) (B), and neutrophil ROS production (log₁₀ MFI) (C). Box plots show distribution with line representing median. Data represent 3 independent experiments with 2 technical replicates per experiment and ≥2 microscopy fields per condition. On ROS plot (B), samples were separated to evaluate the soluble (clarified) and debris compartments. Two-way ANOVA with Tukey’s post-hoc test; *p<0.05, ns=not significant.

### Despite increase ROS production apically with RV replication, there is no impact on RV-A infectious units after incubation with apical neutrophils

Given the findings of increased ROS detected apically in RV infected wells, we wanted to pursue the impact of that ROS on RV infectivity. The impact of ROS on RV’s capsid structure is not known. We assessed viral load by plaque forming units (PFU) and there was no significant decrease in PFU when control or RV-A infected cultures were incubated with apical neutrophils at the same abundance matched to ROS experiments. As previous work in the lab has demonstrated greater virus concentration associated with cellular debris, we chose to assess these fractions separately. We did show increased RV-A infectivity associated with cellular debris compared to free virus (clarified sample), however there was no statistically significant difference in either free or debris associated virus when comparing control or neutrophil incubated hBEC cultures (Fig 5).

**Figure 5:**
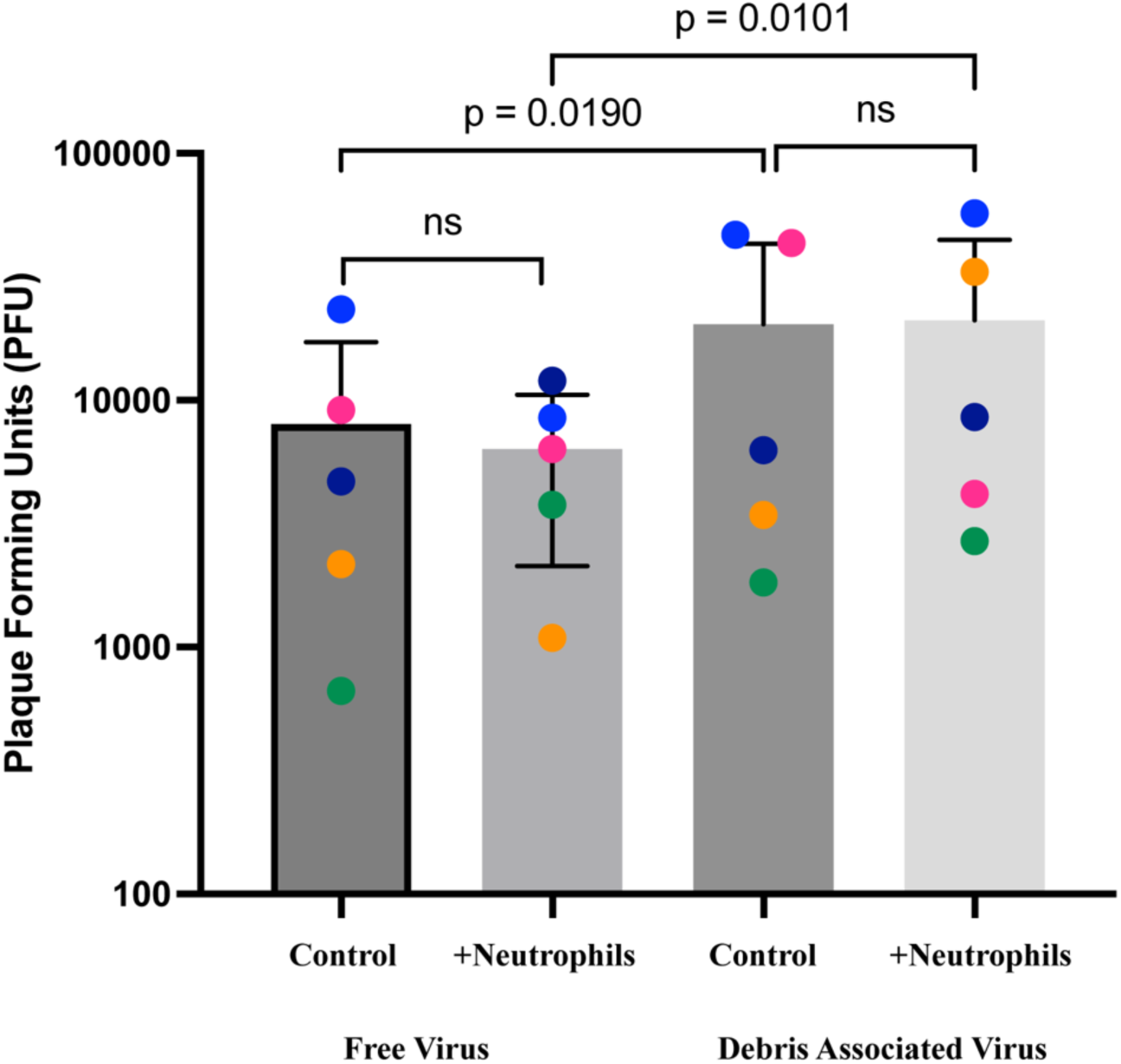
Apical neutrophils do not affect RV-A infectivity after 12 hours; infectious virus is enriched in the debris associated fractions. RV-A infected hBEC donors (n=5, depicted by color) were infected with RV-A for 72 HPI and then incubated with apical neutrophils. Apical washes were then assessed for PFU in both the free virus fraction (clarified) or debris associated fraction. Two-way ANOVA with Tukey’s post-hoc test, *p<0.05, ns=not significant.

## Discussion

Infections with RV and other respiratory viruses strongly induce airway neutrophilia. However, mechanisms of neutrophil activation in the airways and how they contribute to antiviral responses and respiratory symptoms are not well understood. To address this research gap, we used an in vitro model, live cell imaging, and microfluidics to determine how RV infection, both directly to the virus and indirectly through responses of infected bronchial epithelial cells, activates neutrophils. The main findings of our study indicate that there is no direct activation of neutrophils by RV alone. Instead, neutrophil activation arises from polarized epithelial responses: basolateral supernatants from infected epithelial cells increase neutrophil velocity without inducing ROS, whereas apical cellular debris after RV infection does not enhance velocity but does trigger robust ROS release. We also show that in the setting of superoxide production in the presence of RV infection that we do not see an impact on RV infectivity in the presence of neutrophils. These observations identify specific epithelial-neutrophil interactions during RV infection that provides a framework for understanding in vivo post-infection airway neutrophilia.

To understand why RV fails to directly activate neutrophils, we first considered how other common respiratory viruses interact with these cells. Viruses such as influenza have shown direct interaction with neutrophils by binding to sialic acid.(26) Additionally, during acute RSV infection, viral proteins are detected within neutrophils.(27) In fact, RSV was shown to activate neutrophils,(23) decrease peak viral loads and augment degranulation on airway epithelium.(11) Together, these examples demonstrate that direct viral-neutrophil engagement is possible during some respiratory infections. We therefore considered whether receptor availability could explain the absence of direct activation. Neutrophils express minimal CDHR3, the entry receptor used by RV-C, which likely limits viral engagement.(28) Similarly, although neutrophils express ICAM-1 (receptor for RV-B and most RV-A), our data show that neither RV-A nor RV-B induce MPO production, suggesting that RV receptor-neutrophil interactions are insufficient to drive activation. This is consistent with a previous study showing that coincubation of RV-A16 (which binds to ICAM-1) with neutrophils did not induce degranulation.(29)

Because RV does not directly activate neutrophils, we next examined whether epithelial-derived signals could drive neutrophil activation during infection and whether this activation in turn exacerbates epithelial injury. Deng and co-authors demonstrated that apical neutrophils in RSV-infected epithelium exacerbated epithelial damage compared to RSV infection alone(11). In our RV infection models, neutrophils did not enhance RV-associated reductions in barrier integrity or RV-induced cytotoxicity.

Our data showing neutrophil activation only in the setting of airway epithelium led us to hypothesize that ROS production from neutrophils during RV infection was not a direct effect of virus but either cytokine induced activation or an indirect effect of damaged and dying epithelial cells. Utilizing the µBAT microfluidic system, we assessed neutrophil responses with discrete portions of our infected airway epithelium model. The directionality of these activities fits with known sequential patterns of neutrophil trafficking and ROS release. For example, Wang et al. showed that at low (sub nanomolar) concentrations of fMet-Leu-Phe (fMLP) enhance neutrophil chemotaxis; however, at higher doses, increased intracellular Ca²⁺, degranulation, and superoxide production collectively inhibit chemotaxis.(30) The activity in the basolateral supernatant is likely due to RV-induced cytokines and chemokines. RV infection induces many neutrophil chemokines, including CXCL8 and G-CSF. Our data complement previous reports that IL-17C and the neutrophil chemokine CXCL1 can be preferentially secreted in the basal direction, which could promote recruiting neutrophils from the vasculature. (27) (13) (31)

The potent ROS response visualized in our models was from RV infected apical debris. Cell death occurring in the respiratory epithelium exposes damage-associated molecular patterns (DAMPs) that activate neutrophils. By fractionating apical secretions into shed cells or debris and clarified liquid, we ruled out soluble factors (e.g. ATP, HMGB1, calprotectin)(32) as the source of neutrophil activation. Instead, our data show that infected epithelium-associated signals potently activate neutrophil ROS production. Although we show increase in ROS production, this did not equate into lower viral infectivity. Therefore, RV’s capsid structure is likely resistant to the effects of superoxide.

Our results have several strengths and some limitations that should be considered in interpreting these data. Strengths of our data include physiologically relevant cell assessments using neutrophils incubated with differentiated cultures of bronchial epithelium. Additionally, by assessing multiple time points of infection using live cell imaging, we captured dynamic, time-dependent neutrophil activation. Lastly, we measured directional signaling using novel microfluidic techniques. Our study also has some limitations. A limitation of this study is that airway epithelial and neutrophil donors were not matched, and although experiments incorporated multiple independent biological replicates, the use of a simplified in vitro system may not fully capture the complexity of in vivo epithelial–immune interactions. Our experiments best model early stages of infection, but do not address contributions of other cells (e.g., mononuclear cells) or microbes that may modify antiviral and inflammatory effects during the later stages of infection.

Taken together, our findings support a model in which RV-infected airway epithelium generates two distinct classes of neutrophil-modulating signals. Basolateral secretions, likely containing RV-induced cytokines and chemokines, enhance neutrophil migration and recruitment from the vasculature to the airway surface. Once neutrophils have been recruited to the airway, apically shed epithelial cells and associated debris stimulate neutrophil ROS production. These findings indicate that RV-induced epithelial damage or cognate Pathogen-Associated Molecular Patterns (PAMP) signaling, rather than virus alone, triggers ROS activation. Whether this neutrophilic activation increases the proinflammatory state and therefore contributes to immunopathology, or instead has beneficial effects such as promoting healing and tissue repair, is an area for future research.

## Acknowledgments

The author(s) thank the University of Wisconsin Carbone Cancer Center Flow Cytometry Laboratory, supported by P30 CA014520, for use of its facilities and services

## Clinical Implications

Neutrophil activation during rhinovirus infection is driven by epithelial injury rather than direct viral engagement. Because we know that airway neutrophilia correlates with symptoms and airflow obstruction, targeting epithelial damage associated signaling may offer new strategies to reduce RV-related respiratory morbidity.

## Capsule Summary

Neutrophils are not directly activated by rhinovirus but instead respond to polarized signals from infected airway epithelium. Basolateral chemokines enhance migration, while apical epithelial debris triggers ROS release. These directional epithelial and neutrophil interactions help explain how airway neutrophilia develops and contributes to symptoms during RV infection.

## Abbreviations

ROS: Reactive oxygen species
MPO: Myeloperoxidase
PMA: Phorbol myristate acetate
CD11b: Integrin Alpha M
CD66b: CEACAM8
RV qPCR: Rhinovirus quantitative polymerase chain reaction
DAMPS: damage-associated molecular patterns
HPI: hours post-infection
ALI: air liquid interface
hBEC: human bronchial epithelial cells
IL-18: interleukin-18
CXCR2: CXC motif chemokine receptor 2
IL-33: interleukin-33
NET: neutrophil extracellular traps
RSV: respiratory syncytial virus
RV: rhinovirus
TEER: transepithelial electrical resistance
PAMP: pathogen associated molecular patterns
PFU: Plaque forming units
PFUe: Plaque forming units equivalent

---

**Table S1.** CDHR3 Genotype of Cells and Confirmation of RV Replication.

| Experiment | CDHR3<br>genotype | RV-C15 Attachment Control<br>(PFU equivalents) | RV-C15 Replication at 24hrs<br>(PFU equivalents) |
| --- | --- | --- | --- |
| 1 | AA | $3.0 \times 10^6$ | $3.1 \times 10^8$ |
| 2 | AG | $2.8 \times 10^5$ | $8.6 \times 10^7$ |
| 3 | AG | $1.3 \times 10^5$ | $6.9 \times 10^7$ |
| 4 | AA | $2.9 \times 10^5$ | $3.6 \times 10^8$ |

**Table S2:**

| Different curve for at least one data set | Uninfected | 48 HPI RV-C15 | 72 HPI RV-C15 | PMA stimulated |
| --- | --- | --- | --- | --- |
| Best-fit values |  |  |  |  |
| B0 | 1.13 | 1.00 | 0.89 | 3.18 |
| B1 | 0.01 | 0.01 | 0.02 | 0.02 |
| 95% CI (profile likelihood) |  |  |  |  |
| B0 | 0.56 to 1.70 | -0.33 to 2.33 | -0.28 to 2.1 | 1.90 to 4.51 |
| B1 | 0.004 to 0.01 | 0.01 to 0.02 | 0.01 to 0.02 | 0.01 to 0.02 |
| Goodness of Fit |  |  |  |  |
| Degrees of Freedom | 73 | 73 | 73 | 73 |
| R squared | 0.49 | 0.48 | 0.62 | 0.54 |
| Sum of Squares | 117.4 | 646.8 | 503.2 | 643.4 |
| Sy.x | 1.27 | 2.98 | 2.63 | 2.97 |

**Supplemental Figure 1.**
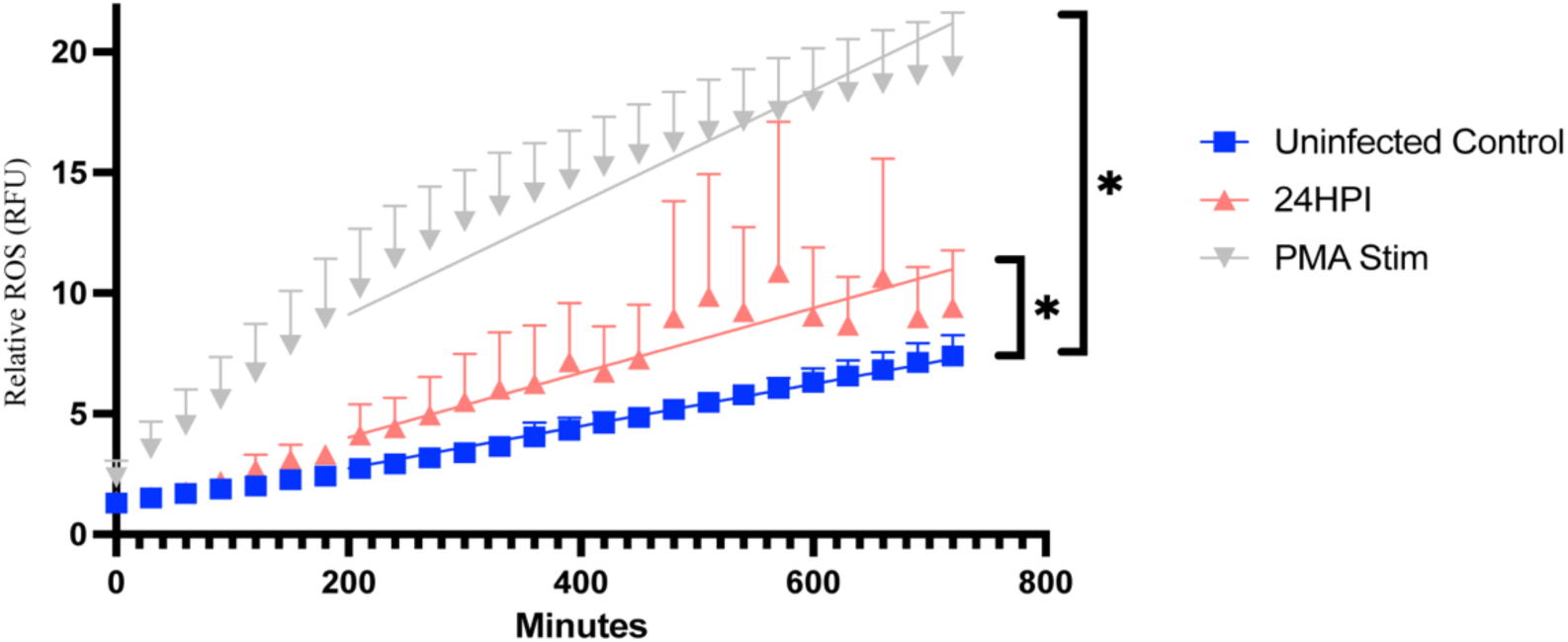
Early neutrophil ROS responses at 24 hours post-infection. hBECs were inoculated with RV-C15 (1×10⁶ PFUe/well) or medium alone and incubated for 24 hours before neutrophils (1×10⁶/well) were added in the presence of CellROX™ Deep Red. Co-cultures were imaged every 30 minutes for 12 hours, and ROS values were normalized to baseline fluorescence. Controls included PMA-stimulated cells (5 ng/mL), uninfected cells, and RV-infected hBECs without neutrophils. Linear regression analysis showed a significant but more variable increase in ROS at 24 HPI. Data represent three independent biological replicates with two technical replicates per condition.

**Supplemental Figure 2:**
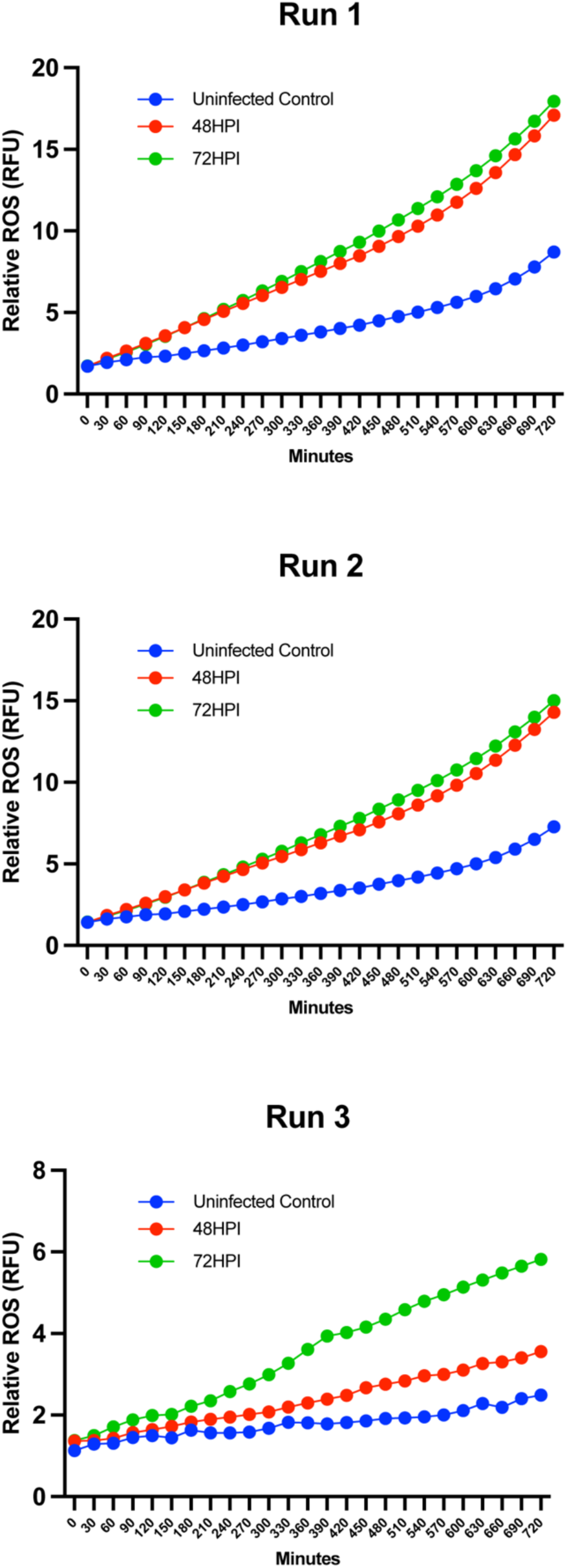
Individual live cell imaging runs. hBECs were inoculated with RV-C15 or medium alone and allowed to replicate for 48 or 72 hours, at which point neutrophils (1×10^6^/well) were added to the apical surfaces of the cell layers in the presence of CellROX™ Deep Red. Data for three independent biological replicates are shown (blue = uninfected control, red= 48 HPI, green = 72 HPI).

**Supplemental Figure 3.**
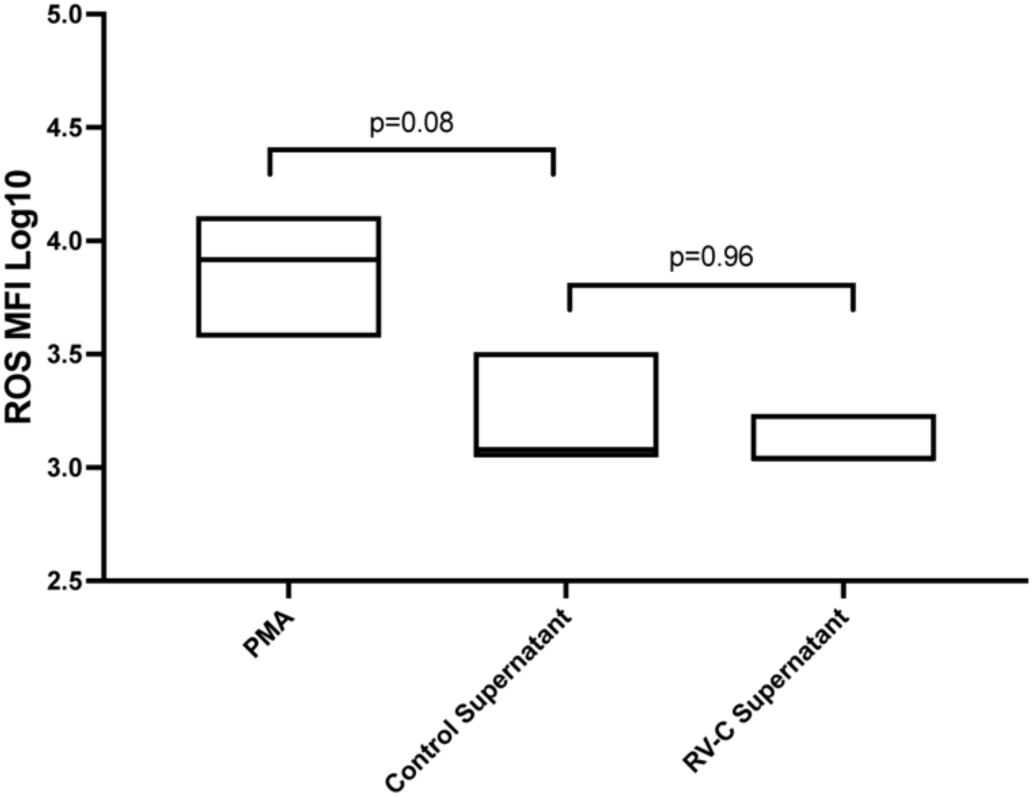
Basolateral supernatants do not induce neutrophil ROS production. Basolateral supernatants collected from medium control- or RV-C15–infected hBECs (72 hours) were applied to freshly isolated neutrophils and ROS production was measured. Neither control nor RV-infected basolateral fractions increased neutrophil ROS relative to control. Box plots show individual measurements with median line. Data represent 3 independent experiments with 2 technical replicates per experiment and ≥2 microscopy fields per condition. Two-way ANOVA with Tukey’s post-hoc test; ns = not significant.

## Declaration of generative AI and AI-assisted technologies in the manuscript preparation process

During the preparation of this work, Microsoft Copilot was used to assist with minor editing and formatting. After using this tool, the author(s) reviewed and edited the content and take full responsibility of the published article.

